# Drought restricted sucrose transport from outer cottonseed coat to fiber and further inhibited cellulose synthesis during cotton fiber thickening

**DOI:** 10.1101/2021.09.14.460198

**Authors:** Honghai Zhu, Wei Hu, Yuxia Li, Jie Zou, Jiaqi He, Youhua Wang, Yali Meng, Binglin Chen, Wenqing Zhao, Shanshan Wang, Zhiguo Zhou

## Abstract

The formation of cotton fiber strength largely relies on continuous and steady sucrose supply to cellulose synthesis and is greatly impaired by drought. However, the effects of drought on sucrose import into fiber and its involvement in cellulose biosynthesis within fiber remain unclear. To end this, moisture deficiency experiments were conducted using two *Gossypium hirsutum* cultivars of Dexiamian 1 (drought-tolerant) and Yuzaomian 9110 (drought-sensitive). Fiber strength was significantly decreased under drought. The results of ^13^C isotope labeling indicated that drought notably reduced sucrose efflux from cottonseed coat to fiber, and this was caused by down-regulation of sucrose transporter genes (*GhSWEET10* and *GhSWEET15*) in the outer cottonseed coat, finally leading to decreased sucrose accumulation in fiber. Further, under drought, the balance of sucrose allocation within fiber was disrupted by increasing the flow of sucrose into β-1,3-glucan synthesis and lignin synthesis but hindering that into cellulose synthesis in both cultivars. Additionally, glycolysis and starch synthesis were specifically enhanced by drought in Yuzaomian 9110, which further reduced the flow of sucrose into cellulose synthesis. Under drought, the cellulose deposition was decreased due to promoted cellulose degrading process in Dexiamian 1 and stunted cellulose synthesis in Yuzaomian 9110. Consequently, reduced cellulose content was measured in drought-stressed fibers for both cultivars. In summary, the inhibited cellulose accumulation caused by drought was mainly due to reduced sucrose translocation from the outer cottonseed coat to fiber, and less sucrose partitioned to cellulose synthesis pathway under the condition of intensified competition for sucrose by different metabolic pathways within fiber, finally degrading the fiber strength.

**Highlight:** This article revealed the path of sucrose flow from cottonseed coat to cotton fiber and sucrose competition patterns within cotton fiber under drought and their relationships with fiber strength loss.

## Introduction

Upland cotton (*Gossypium hirsutum*), one of the most important industrial crops in the world (Ruan, 2013), produces up to 95% of the natural lint fiber used for the textile industry (Huang *et al*., 2021). Although cotton is typically grown in heat and semi-arid regions, it is sensitive to drought (Viglioni *et al*., 1998), especially during the flowering and boll-forming stage when the occurrence of drought could do great damage to the fiber yield and quality (Snowden *et al*., 2014). Over the last decades, drought has caused dramatic losses in cotton yield and quality every year (Abdelraheem *et al*., 2019). Fiber strength is a key parameter for fiber quality assessment and is closely related to yarn property. Mild drought has little effect on fiber strength, but as severe drought occurred, fiber strength was dramatically reduced (Basal *et al*., 2009; Dağdelen *et al*., 2009; Wang *et al*., 2016; Hu *et al*., 2018). Climate change scenarios are predicting that the frequency, severity, and duration of drought will significantly increase in the near future (Dai, 2011), posing a bigger threat to cotton fiber development and fiber strength.

Cotton fiber development is identified as a progressive process of cellulose deposition and cell wall construction. At about 16 days post anthesis (DPA), the cotton fiber enters into the secondary wall synthesis stage which ceases at around 40 DPA (Haigler *et al*., 2012). The formation of cotton fiber strength largely depends on the cellulose synthesis and deposition during this phase (Gou *et al*., 2007), as cellulose accounts for up to 95% of secondary wall dry weight at maturity (Haigler *et al*., 2012). Cellulose biosynthesis is located at the plasmamembrane and requires the participation of a variety of enzymes and substances. Sucrose is widely believed to be the initial precursor for cellulose synthesis and it has to be degraded into uridine-5’-diphosphoglucose (UDPG) by sucrose synthase (SuSy) before streaming into cellulose biosynthesis pathway (Carpita and Delmer, 1981; Coleman *et al*., 2009). Then the cellulose synthase (CesA) complexes, also known as the rosettes, catalyze UDPG to produce β-(1,4)-linked glucan chains (Read and Bacic, 2002) that are organized into cellulose. Except for cellulose biosynthesis, the imported sucrose is also consumed by many other metabolic pathways like respiration and biosynthesis of β-1, 3-glucan (callose), lignin, and starch (Amthor, 2003; Koch, 2004; Scheible and Pauly, 2004; Farrokhi *et al*., 2006). These metabolisms could affect cellulose synthesis and cell wall formation by competing for sucrose. For instance, inhibited cellulose biosynthesis was observed when starch biosynthesis was enhanced in some mutants of *Pisum sativum* and *Arabidopsis thaliana* (Harrison *et al*., 1998; Peng *et al*., 2000), and the down-regulation of lignin biosynthesis is accompanied by an increase in cellulose biosynthesis in *Populus tremuloides* mutants (Hu *et al*., 1999). It has been reported that drought decreased cellulose biosynthesis in cotton fiber (Ibrahim *et al*., 2019; Gao *et al*., 2020). Unfortunately, whether the reduction of cellulose under drought is somehow related to changes in biosynthesis of β-1, 3-glucan, lignin, or starch, etc. is not clear, which hinders our panoramic understanding of cotton fiber development under drought.

The developing cottonseed can be divided into three parts: the fiber, also known as trichome to the cottonseed coat epidermis, the cottonseed coat where phloem terminates and the embryo predominately consisting of cotyledons (Hendrix, 1990; Ruan *et al*., 1997). Since the fiber initiates from the cottonseed coat, its development is tightly controlled by the cottonseed coat which works as a nutrients transfer station (Ruan and Furbank, 2003; Ruan, 2013). As the principal form of photosynthates in higher plants, sucrose is primarily produced in the photosynthetic organs and transported to the sink via phloem (De Schepper *et al*., 2013). The sucrose is firstly unloaded from the phloem in the outer cottonseed coat and then transported outwards to the fiber (Ruan *et al*., 1997; Ruan and Chourey, 1998). The transfer efficiency of sucrose in cottonseed coat is an essential limiting factor for the development of cottonseed and fiber (Pugh *et al*., 2010). The sucrose transporters: SUT (sucrose transporter) also called SUC (sucrose carrier), and SWEET (sugars will eventually be exported transporter) are believed to participate in the sucrose transport and movement in plants (Ayre, 2011; Zhang and Turgeon, 2018). Sucrose-specific SWEETs mediate passive sugar efflux out of the cytosol and SUTs catalyze active uptake of sucrose from the apoplast against concentration gradient energized by ATP (Baker *et al*., 2012; Chen *et al*., 2015). Both transport processes are involved in phloem loading in source leaves as well as in the post-phloem pathway in sink tissues (Chen *et al*., 2015). Nine SUTs were identified in *Arabidopsis* and classified into three types (Kühn and Grof, 2010; Peng *et al*., 2020). On this basis, a total of nine pairs of homologous SUT genes were identified in *Gossypium hirsutum* and they participated in the responses to multiple abiotic stresses (Li *et al*., 2018). *GhSWEET10, GhSWEET12*, and *GhSWEET15* have been demonstrated to mediate sucrose efflux and involve fiber development and biotic stress in cotton (Cox *et al*., 2017; Sun *et al*., 2019; Ding *et al*., 2021). Nonetheless, little is known about sucrose translocation from the cottonseed coat to the fiber under drought and its relationship with fiber strength loss needs to be elucidated.

Therefore, it was hypothesized that (1) drought would alter the efficiency of sucrose transfer from cottonseed coat to fibers, and (2) drought would further inhibit sucrose flow to the cellulose synthesis pathway. In this study, the objectives were 1) to explore the effects of drought on sucrose import into fibers, and 2) to elucidate the effects of drought on the flow of sucrose into different metabolic pathways within fibers. The results will broaden our understanding of fiber strength decline under drought conditions, and provide new ideas for breeding drought-resistant varieties and improving fiber quality.

## Materials and methods

### Experimental design and treatment

The experiments were carried out in an open-top rain-proof shed (25 m long, 6 m wide, 3 m high) at Pailou experimental station of Nanjing Agricultural University, Nanjing (118.78°E, 32.04 °N), Jiangsu, China from 2018 to 2019. Two *Gossypium hirsutum* cultivars, Dexiamian 1 (drought-tolerant) and Yuzaomian 9110 (drought-sensitive) (Zou *et al*., 2020), were selected as plant materials. Cottonseeds were sown in nutrimental bowls on 7 April 2018, and 12 April 2019, respectively. As the third true leaves were fully expanded, thriving seedlings were selected and transplanted into plastic pots containing around 12 kg (dry weight) soil (clay, mixed, thermic, Typic Alfisols). Each pot contained one cotton seedling. Before different water regimes were established, equivalent and sufficient irrigation was applied to every plant to keep the soil relative water content (SRWC) at (75±5)% which was the optimum soil moisture for cotton growth in this type of soil (Wang *et al*., 2016).

When approximately 50% of white flowers at the first fruit nodes of the middle fruit branches (4-6 fruit branches) bloomed, two levels of SRWC (75±5)% (control, CK) and (45±5)% which was considered severe drought that significantly impaired the fiber yield and quality in this type of soil (Wang *et al*., 2016), were administered until boll opening. During the trial period, soil samples were taken every 2-3 days and dried in a 105°C ventilated oven for 8 hours to estimate SRWC gravimetrically according to Wang *et al*. (2016).

### Sampling method

White flowers at the first fruit nodes of the middle fruit branches were tagged and marked as 0 days post anthesis (DPA). Six to eight tagged bolls were harvested every seven days at 9:00 am from 17 DPA to 38 DPA. The cottonseeds were quickly stripped from the boll and were separated into fibers, cottonseed coats, and embryos. In addition, the outer cottonseed coat was further isolated at 17 and 24 DPA. All samples were immediately immersed into liquid nitrogen and stored at -80°C for subsequent physiological and molecular determinations.

### Fiber yield and quality

Tagged mature cotton bolls of 20 from each treatment were reaped and air-dried. Each boll was weighed and ginned individually, then the seeds and lint were weighed separately. Moreover, seed number in each boll was counted. The lint index (fiber weight per 100 seeds) was calculated. Finally, fiber strength was measured with six replications by using Uster HVI MF100 (USTER®, Uster, Zurich, Switzerland).

### Biomass accumulation and distribution

For cotton boll biomass accumulation, three tagged bolls per treatment at 17, 24, 31, 38 DPA and maturation were harvested. Fibers and cottonseed coats were isolated from ten seeds of each boll, weighed for the fresh weight (FW), dried at 80°C to constant weight for the dry weight (DW), and biomass per seed was calculated.

The biomass distribution was calculated as follows:

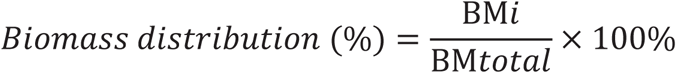

Where BM*i* represents the biomass of fibers, cottonseed coats, or embryos of ten seeds and the sum-up of the three parts makes BM*total* value.

### Cottonseed coat observation and measurement of cottonseed coat thickness

For cottonseed coat observation, thin slices were cross-cut from fresh seeds in the middle position with a sharp razor blade. Slices were examined on a stereomicroscope (SZX16, Olympus Corporation, Japan). Digital images were captured and stored as TIFF files and the thickness of cottonseed coat, inner cottonseed coat, and outer cottonseed coat was measured by using the Image-pro Plus program (Olympus).

### ^13^CO_2_ feeding experiment and ^13^C-photoassimilate translocation rate within the cottonseeds

To investigate the photosynthates translocation rate within cottonseeds, the ^13^CO_2_ feeding experiment was conducted in 2019. Because both Dexiamian 1 and Yuzaomian 9110 are short-season cotton cultivars and their bolls start to open at about 38 DPA, ^13^C-photoassimilate translocation rate within cottonseeds was measured at 17, 24, and 31 DPA. The isotope labeling method was based on the report of Hu *et al*. (2020). Three bolls from each treatment were marked with plastic tags, and each leave subtending the boll was placed into a sealed transparent chamber with pre-injected 5 ml of ^13^CO_2_ (Shanghai Research Institute of Chemical Industry, China) for four hours from 08:30 h -12:30 h. After 24 hours at 8:30 h, the marked bolls were harvested and swiftly divided into fibers, cottonseed coats, and embryos. Then they were dried in a ventilated oven at 80°C to constant weight and were grounded into fine powder for the following tests.

About 3 mg of finely grounded sample powder was used to detect the total carbon and the ^13^C abundance (δ^13^C), with an EA-1110 elemental analyzer (Carlo Erba Thermoquest, Milan, Italy) at 1020 °C coupled to an isotope ratio mass spectrometer (Finnigan MAT, Bremen, Germany). The standard was applied according to the international Pee Dee Belemnite standard (Pee-Dee Belemnite, SC, USA). The ^13^C distribution ratio among each tissue was calculated according to Ruehr *et al*. (2009).

^13^C content in a given sample was calculated as follow:

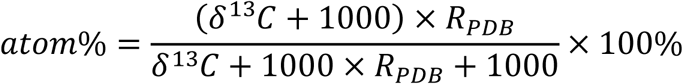

Where atom% is the ^13^C abundance of the labeled tissue, R is the corresponding ratio ^13^C/^12^C, the δ value represents the ratio of heavy and light isotopes in the samples compared with the standard reference material.

The ^13^C accumulation in the given tissue was calculated as follow:

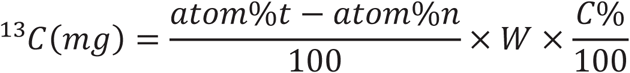

Where atom%t is the atom% of ^13^C of labeled tissue, atom%n is the atom% of ^13^C of unlabeled natural tissue, W is the dry weight of each tissue (mg), C% is the percentage of total carbon content in each tissue.

The ^13^C distribution ratio was calculated as:

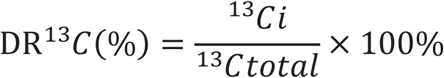

Where ^13^Ci represents the biomass of fibers, cottonseed coats, or embryos per boll and the sum-up of the three parts makes ^13^Ctotal value.

### Determination of the content of sucrose, glucose, fructose, cellulose, β-1, 3-glucan, and starch

Soluble sugars were extracted and determined according to Hu *et al*. (2020) with slight modification. Frozen samples of 0.2 g were grounded into fine powder with liquid N_2_, then transferred into tubes with 80% ethanol, extracted at 80°C for three times, and after centrifugation, the supernatants were combined and diluted to 25 ml with 80% ethanol. Extraction of 20 μL was pipetted into a colorimetric 96-well microtiter plate. After drying at 45°C, 20 μL of distilled water was added into each well and let it stand for 10 min. For the determination of glucose, 100 μL of glucose assay reagent (Sigma-Aldrich) was added to each sample followed by incubation at 30°C for 15 min. To measure the fructose, the samples were premixed with 0.25 U phosphoglucose isomerase and then incubated at 30°C for 15 min. For the measurement of sucrose, 83 U invertase was added into each sample and the mixture was incubated at 30°C for 60 min. The absorbances of the mixtures at 340 nm wavelength were read after each incubation.

Starch was extracted and determined according to Hu *et al*. (2018) with some modifications. Dried cotton fiber of 0.2 g was washed with distilled water for three times to remove soluble sugars. The residues were digested with 2 mL of 1M KOH at 100°C for 60min. Next, the pH of extraction was adjusted to 6.5–7.5 with 0.2 M acetic acid. After that, 250 μL of amyloglucosidase was added to the mixture followed by incubation at 65°C for 60 min to convert starch into glucose. Finally, the glucose content in the mixture was determined according to the aforementioned method.

Cellulose content was measured with the anthrone method (Updegraff, 1969). In brief, 0.2 g dried fiber was immersed into the acetic–nitric acid reagent for digestion, washed with distilled water, and dried again. Then the samples were dissolved in 67% H_2_SO_4_ and added with 0.2% anthrone reagent for reaction. The absorbance of the reaction mixture was measured using a UV–Vis spectrophotometer at 625 nm and the calibration was prepared with microcrystalline cellulose. β-1, 3-glucan content was determined using a modified method described by (Köhle *et al*., 1985). Dried cotton fiber of 0.2 g was preprocessed in 10 ml of 80% ethanol containing 10 mM EDTA for 30 min to remove autofluorescent soluble compounds. After dried, the samples were soaked with 10 ml of 1M NaOH, incubated at 80°C for 30 min to solubilize the β-1, 3-glucan, and centrifuged for 15 min at 380 g. Next, 600 µl of supernatant was mixed with 0.1% aniline blue solution (400 µl), 1 M HCl (210 µl), and 1 M glycine/NaOH buffer (590 µl, pH 9.5). Then the mixture was water bathed at 50°C for 20 min and cooled at room temperature for 30 min. The fluorescence was read with a fluorimeter using 400 nm excitation light and 510 nm emission light. The β-1, 3-glucan content was calculated according to the calibration curve established by using a freshly prepared solution of pachyman (β-1, 3-glucan) in 1 M NaOH.

### Transcriptome analysis

The fiber from CK and drought treatments were sampled with three biological replicates at 17 DPA in 2019. The total RNA of each sample was individually extracted using RNAprep Pure Plant Kit (Polysaccharides & Polyphenolics-Rich) (Vazym, China). Following RNA-sequencing (RNA-seq) analysis based on next-generation sequencing and estimation of transcript expression levels were conducted by Allwegene (Beijing, China). After RNA libraries were produced, they were sequenced using Illumina second-generation high-throughput sequencing platform (HiSeqTM 2500/4000) and PE150 sequencing strategy. Attained clean reads were assembled to *Gossypium hirsutum* genome (*Gossypium hirsutum*, CRI; https://cottonfgd.org/about/download/assembly/genome.Ghir.CRI.fa.gz) using Hisat2 v2.0.4. The gene expression levels were normalized and computed as transcripts per million (TPM) (Vera *et al*., 2019). For differential expression analysis, genes with a |log_2_ fold change|>1 and an adjusted *P*-value or false discovery rate (FDR)<0.05 were designated as differentially expressed genes (DEGs). Gene function annotation was defined by using the *Gossypium hirsutum*, CRI database (https://cottonfgd.org/about/download/annotation/gene.Ghir.CRI.gff3.gz). Analysis of gene ontology (GO) term enrichment and pathway enrichment was conducted using GOseq (Young *et al*., 2010) and Kyoto Encyclopedia of Genes and Genomes (KEGG) (Kanehisa *et al*., 2008) database respectively. GO terms or pathways with a *P*-value≤0.05 were considered as differentially regulated.

### Quantitative real-time PCR (qRT-PCR) analysis

Total RNA of the outer cottonseed coat and the fiber was extracted using RNAprep Pure Plant Kit (Polysaccharides & Polyphenolics-Rich) (Vazym, China). Then total RNA of 1 μg was used to produce cDNA with PrimeScript™ RT Master Mix (Vazym, China). qRT-PCR was performed using TB Green™ Premix Ex Taq™ II (Vazym, China) on a CFX Connect (TM) Real-Time PCR Detection System (BIO-RAD, Singapore). The results obtained were standardized to the expressions of a cotton polyubiquitin gene (RUBIQ1) and gene expression levels were calculated according to 2^-ΔCT^ method in the outer cottonseed coat and 2^-ΔΔCT^ method in the fiber, respectively. All primers used in this study are listed in Table S1.

### Statistical analysis

The data presented in tables and figures are the mean ± standard deviation (SD) of at least three replications. The effects of water treatment, cultivar, and the interaction of the two factors on lint index, fiber strength, and biomass of fiber, cottonseed coat, and embryo within a given year were assessed using a two-way analysis of variance (ANOVA) and the significant difference was determined with the least significant difference (LSD) test (*P*<0.05). The other parameters with the same cultivar between water treatments within a given year were processed using a one-way analysis of variance (ANOVA) and the comparison of means was performed using Student’s *t*-test (*P*<0.05). All statistical analyses were performed using SPSS 26.0 software. Figures were generated by using Origin 2021 software.

## Results

### Effects of drought on cotton lint index and fiber strength

Under drought, lint index and fiber strength were dramatically decreased by 4.6-10.5% and 4.8-5.8% in Dexiamian 1, and 11.6-16.1% and 9.4-10.8% in Yuzaomian 9110, respectively (Table. 1). In addition, a significant difference in the interaction effect of water treatment×cultivar was observed in both 2018 and 2019.

**Table 1.** Effects of drought on lint index and fiber strength in 2018 and 2019

| Cultivar | Water treatment<br>(WT) | Lint index (g) |  | Fiber strength (cN tex<sup>-1</sup>) |  |
| --- | --- | --- | --- | --- | --- |
|  |  | 2018 | 2019 | 2018 | 2019 |
| Dexiamian 1 | CK | 6.57±0.19a | 6.96±0.34a | 28.76±0.55a | 28.42±0.65a |
|  | Drought | 6.27±0.25a | 6.23±0.32b | 27.13±0.65b | 27.07±0.53b |
| Yuzaomian 9110 | CK | 7.57±0.35a | 7.44±0.31a | 30.55±0.66a | 29.97±1.02a |
|  | Drought | 6.35±0.25b | 6.58±0.24b | 27.69±0.57b | 26.72±0.66b |
| Significance of factors |  |  |  |  |  |
| WT |  | ** | ** | ** | ** |
| Cultivar |  | ** | ** | * | ns |
| WT×Cultivar |  | ** | ns | * | * |

### Effects of drought on biomass distribution within cottonseed

Under drought, the biomass of fiber and embryo at maturity stage was significantly decreased, and the biomass of cottonseed coat was little affected (Table S2). Drought notably enhanced the proportion of biomass partitioned to cottonseed coat, but decreased the proportion distributed to fiber, and the fraction of embryo was not affected (except for Yuzaomian in 2019, Fig. 1A). Under drought, the biomass accumulation of cottonseed coat was elevated during 17-24 DPA (Fig. 1B), and that of fiber was significantly decreased, especially during 17-31 DPA (Fig. S1). As presented in Fig. 1C, the cottonseed coat can be clearly identified into two parts: outer cottonseed coat and inner cottonseed coat. The thickness of cottonseed coat, outer cottonseed coat, and inner cottonseed coat was measured and was little affected under drought (Fig. S2).

**Fig. 1.**
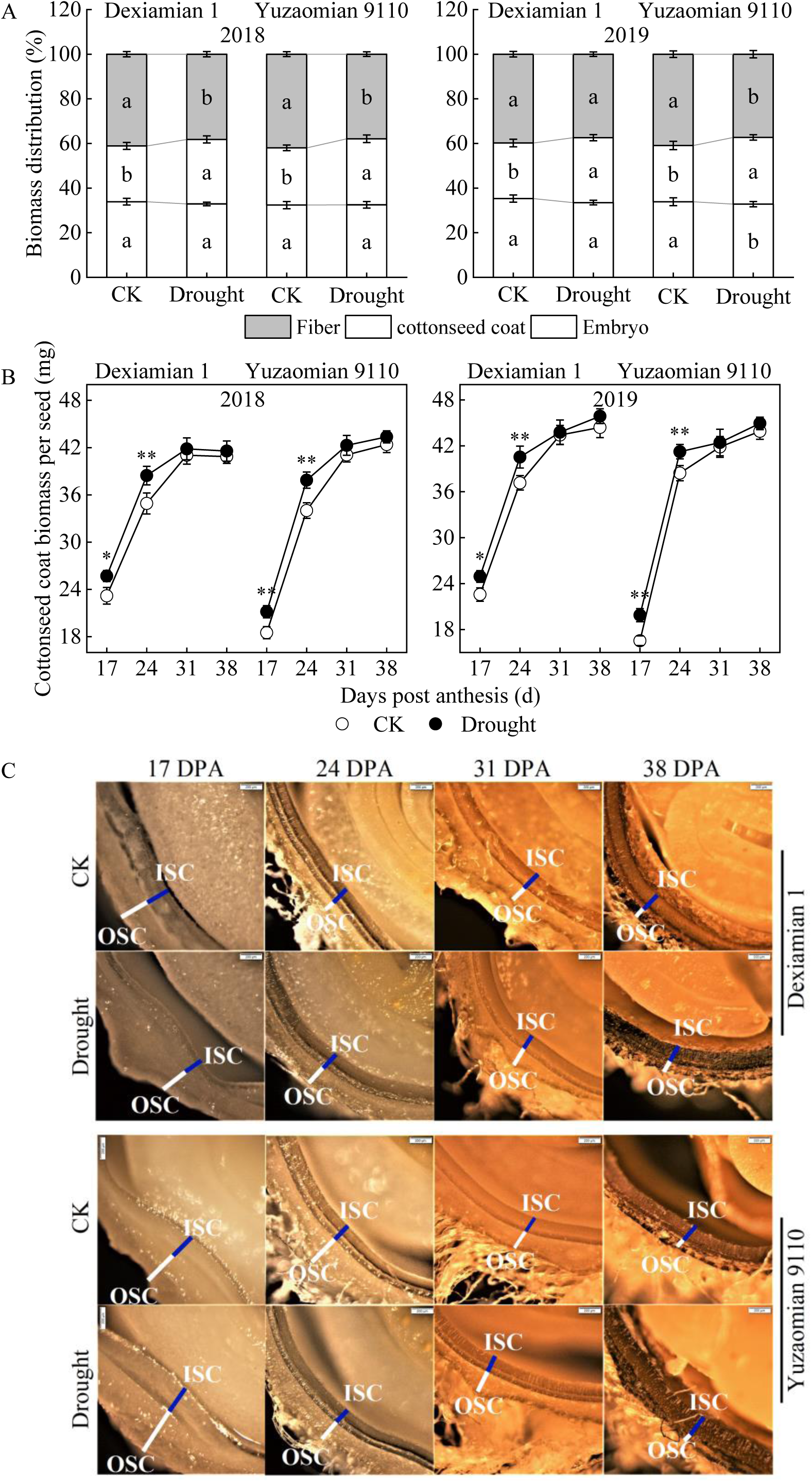
Biomass distribution among embryo, cottonseed coat and fiber at maturity stage (A), dynamic changes of cottonseed coat biomass per seed (B) in 2018 and 2019, and dynamic observation of cottonseed coat slices in 2019 (C) in different treatments. OSC, outer cottonseed coat; ISC, inner cottonseed coat. Scale bars in Fig. 1C: 200 μm. Values are means of three replicates ±SD. Different letters in Fig. 1A indicate statistically significant differences between different treatments (*P*<0.05). * and ** in Fig. 1B indicate statistically significant differences determined between different treatments at *P*=0.05 and *P*=0.01, respectively.

### Effects of drought on newly produced photosynthates translocation to fiber

Under drought, ^13^C distribution ratio (DR_13C_) value in fiber was consistently lower by comparing to CK, and an opposite tendency was observed in cottonseed coat (Fig. 2). Meanwhile, DR_13C_ value in embryo under drought was increased at 17 DPA but decreased during 24-31 DPA (Fig. 2). These results indicated that drought affected the photosynthates transportation and distribution patterns in developing cottonseed.

**Fig. 2.**
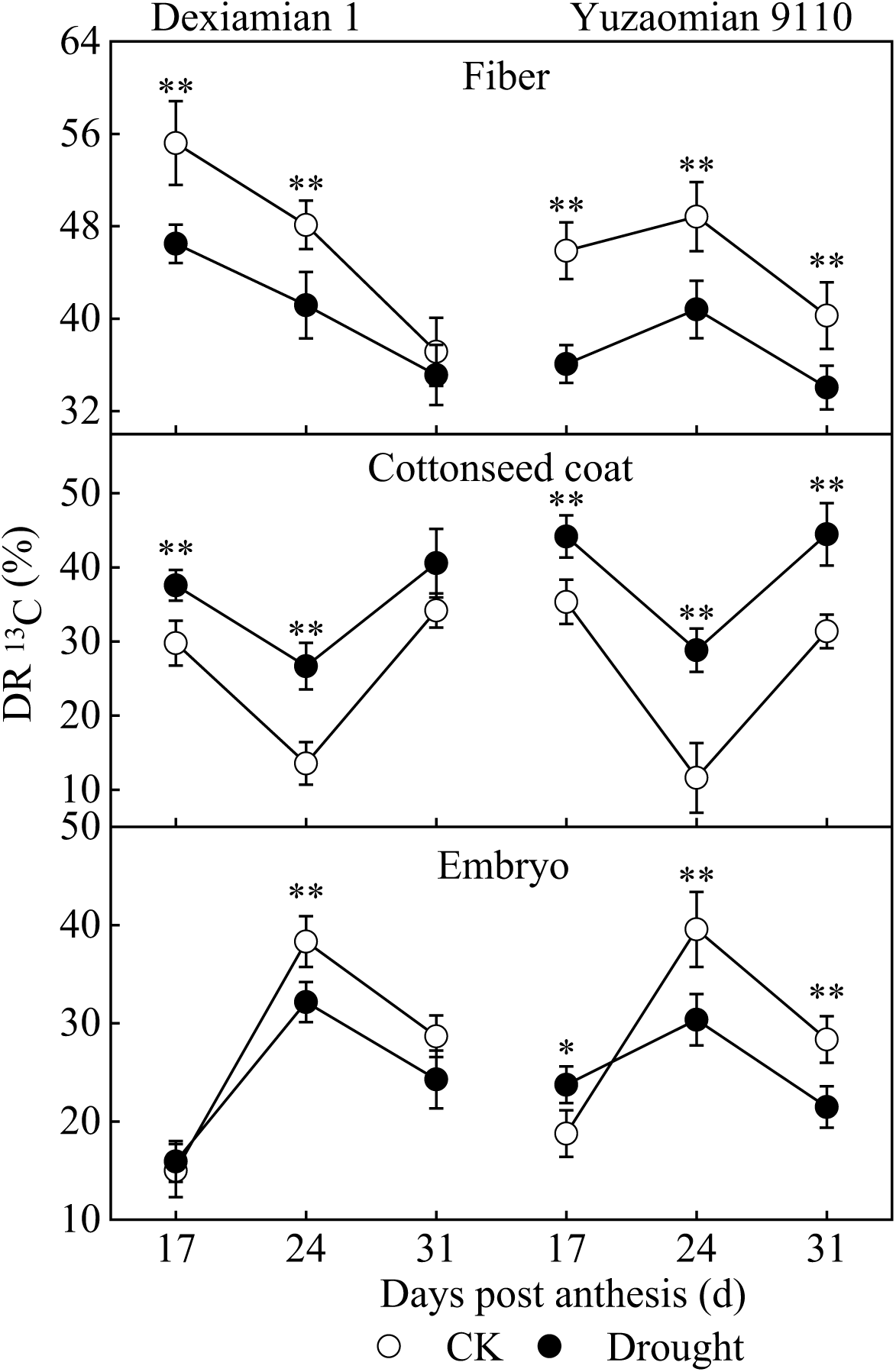
Effects of drought on the relative distribution of ^13^C among fiber, cottonseed coat and embryo in 2019. All values are the mean of three replicates ±SD. * and ** indicate statistically significant differences determined between different treatments at *P*=0.05 and *P*=0.01, respectively.

### Effects of drought on sucrose content in cottonseed coat and fiber

As the main form of photosynthates for transportation, sucrose content in fiber and cottonseed coat was measured. Compared with CK, sucrose level in cottonseed coat was significantly enhanced during 17-24 DPA while was little affected during 31-38 DPA under drought (Fig. 3A). Oppositely, the fiber sucrose content was notably decreased under drought during 17-31 DPA (Fig. 3B).

**Fig. 3.**
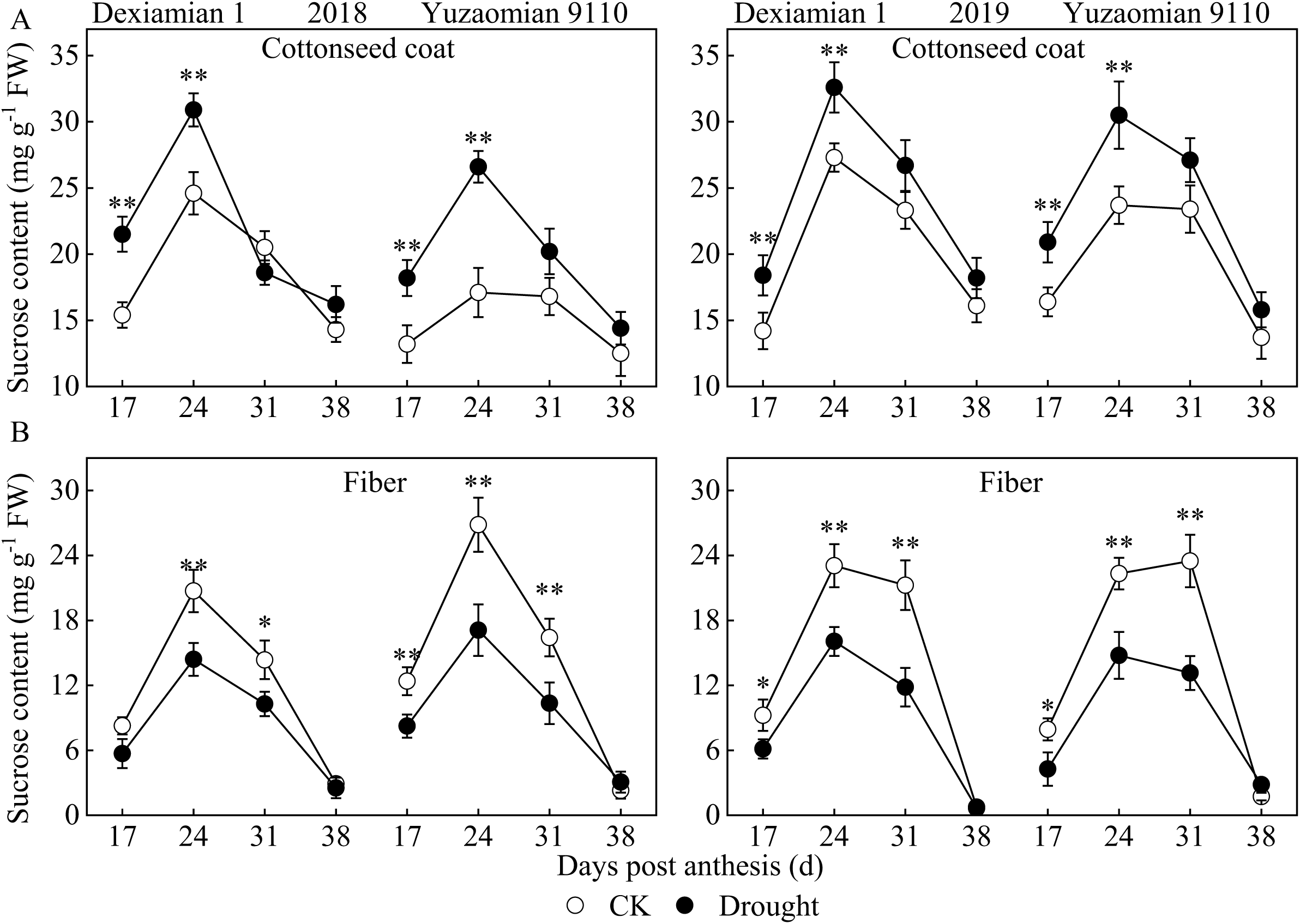
Effects of drought on the sucrose content in cottonseed coat (A) and fiber (B) in 2018 and 2019. Values are the mean of three replicates ±SD. * and ** indicate statistically significant differences determined between different treatments at *P*=0.05 and *P*=0.01, respectively.

### Effects of drought on the expressions of sucrose transporter genes in the outer cottonseed coat

A total of nine pairs of homologous SUT genes have been identified in *Gossypium hirsutum* (Li *et al*., 2018). Among the nine paralogous SUT genes, *GhSUT1A/D, GhSUT3A/D, GhSUT7A/D*, and *GhSUT9A/D* were not detectable in the outer cottonseed coat, thus only *GhSUT2A/D, GhSUT4A/D, GhSUT5A/D, GhSUT6A/D*, and *GhSUT8A/D* were shown in this work (Fig. 4). Overall, the expression levels of *SUTs* (except *GhSUT5A/D*) in the outer cottonseed coat were markedly promoted at 17 DPA under drought, especially *GhSUT2A/D* and *GhSUT4A/D* (Fig. 4). At 24 DPA, the expression levels of *GhSUT4A/D, GhSUT6A/D*, and *GhSUT8A/D* in the outer cottonseed coat were significantly decreased, while the expression of *GhSUT5A/D* showed an opposite changing trend and *GhSUT2A/D* was little affected (Fig. 4). SWEET10, SWEET12, and SWEET15 were confirmed to be sucrose effluxer in cotton (Cox *et al*., 2017; Sun *et al*., 2019; Ding *et al*., 2021). Under drought, *GhSWEET10* and *GhSWEET15* in the outer cottonseed coat were significantly down-regulated at 17 and 24 DPA, and the expression of *GhSWEET12* was little affected at 17 DPA but significantly up-regulated at 24 DPA (Fig. 4).

**Fig. 4.**
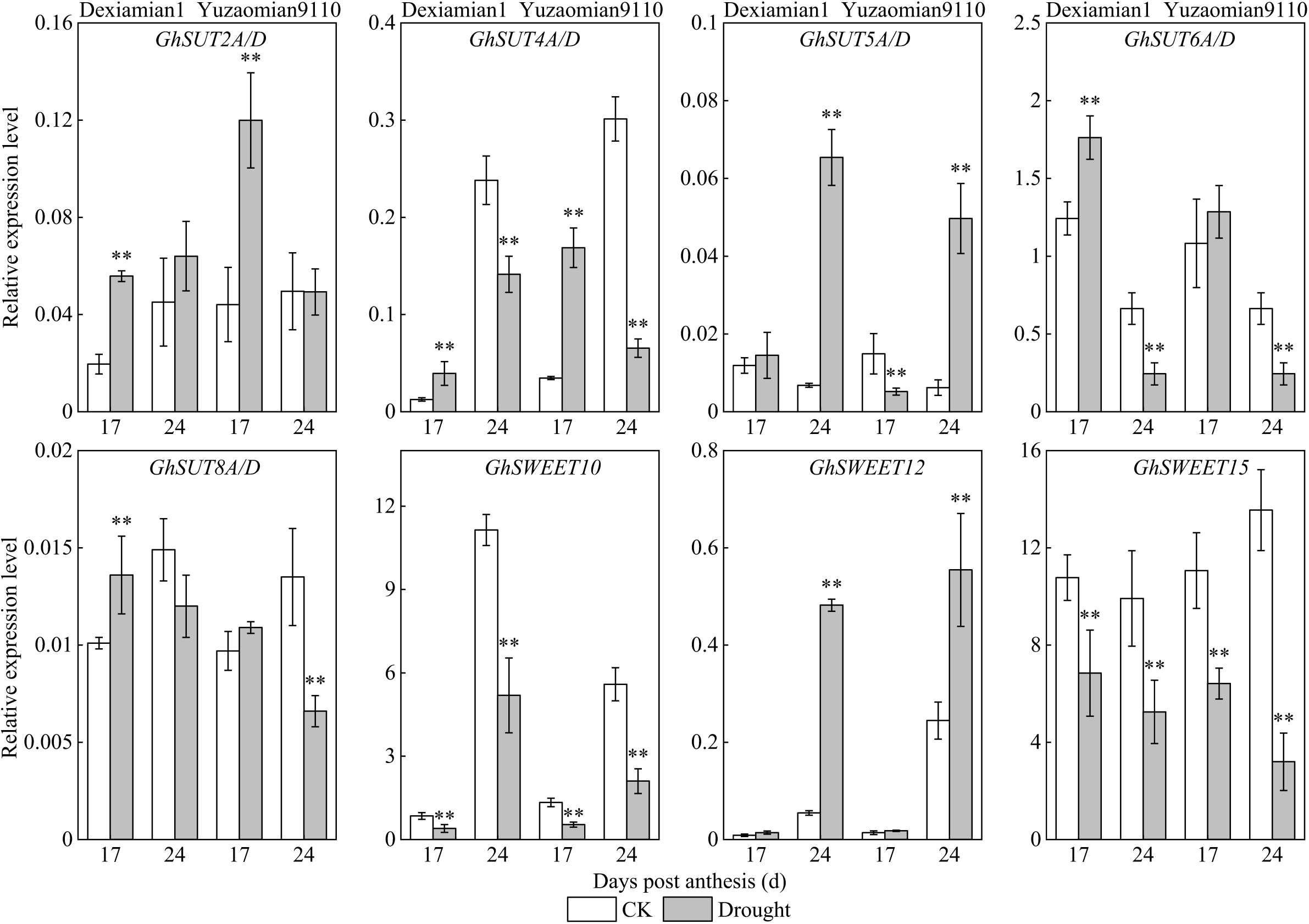
Effects of drought on the relative expression levels of sucrose transporter genes in the outer cottonseed coat at 17 and 24 DPA in 2019. Values are the means of three replicates ±SD. * and ** indicate statistically significant differences determined between different treatments at *P*=0.05 and *P*=0.01, respectively.

### Changes of cellulose, β-1, 3-glucan, and starch contents in cotton fiber

Under drought, the cellulose content in mature cotton fiber was significantly decreased by 5.2-5.6% in Dexiamian 1 and 5.5-6.1% in Yuzaomian 9110, respectively (Table. 2). In developing cotton fiber, the cellulose accumulation was notably decreased under drought (Fig. 5A). While β-1, 3-glucan content was elevated under drought, and the largest gap between drought and CK appeared at 17 DPA (Fig. 5B). Compared with CK, starch content in fiber significantly increased in Yuzaomian 9110 at 17 DPA under drought, and little significant difference was observed in Dexiamian 1 (Fig. 5C).

**Table 2.**
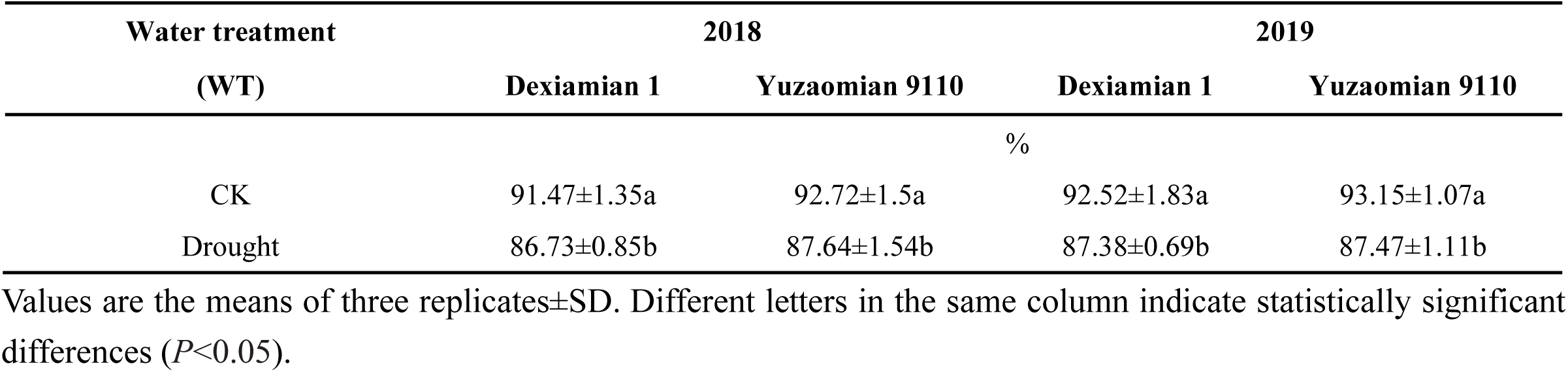
Effects of drought on the cellulose content in mature cotton fiber in 2018 and 2019.Values are the means of six replicates±standard deviation (SD). Different letters within in the same column indicate statistically significant differences (*P*<0.05). * and ** indicate statistically significant differences at *P*=0.05 and *P*=0.01, respectively, and ns means no significant difference.

| Water treatment<br>(WT) | 2018 |  | 2019 |  |
| --- | --- | --- | --- | --- |
|  | Dexiamian 1 | Yuzaomian 9110 | Dexiamian 1 | Yuzaomian 9110 |
| % |  |  |  |  |
| CK | 91.47±1.35a | 92.72±1.5a | 92.52±1.83a | 93.15±1.07a |
| Drought | 86.73±0.85b | 87.64±1.54b | 87.38±0.69b | 87.47±1.11b |

Values are the means of three replicates±SD. Different letters in the same column indicate statistically significant differences ( $P<0.05$ ).

**Fig. 5.**
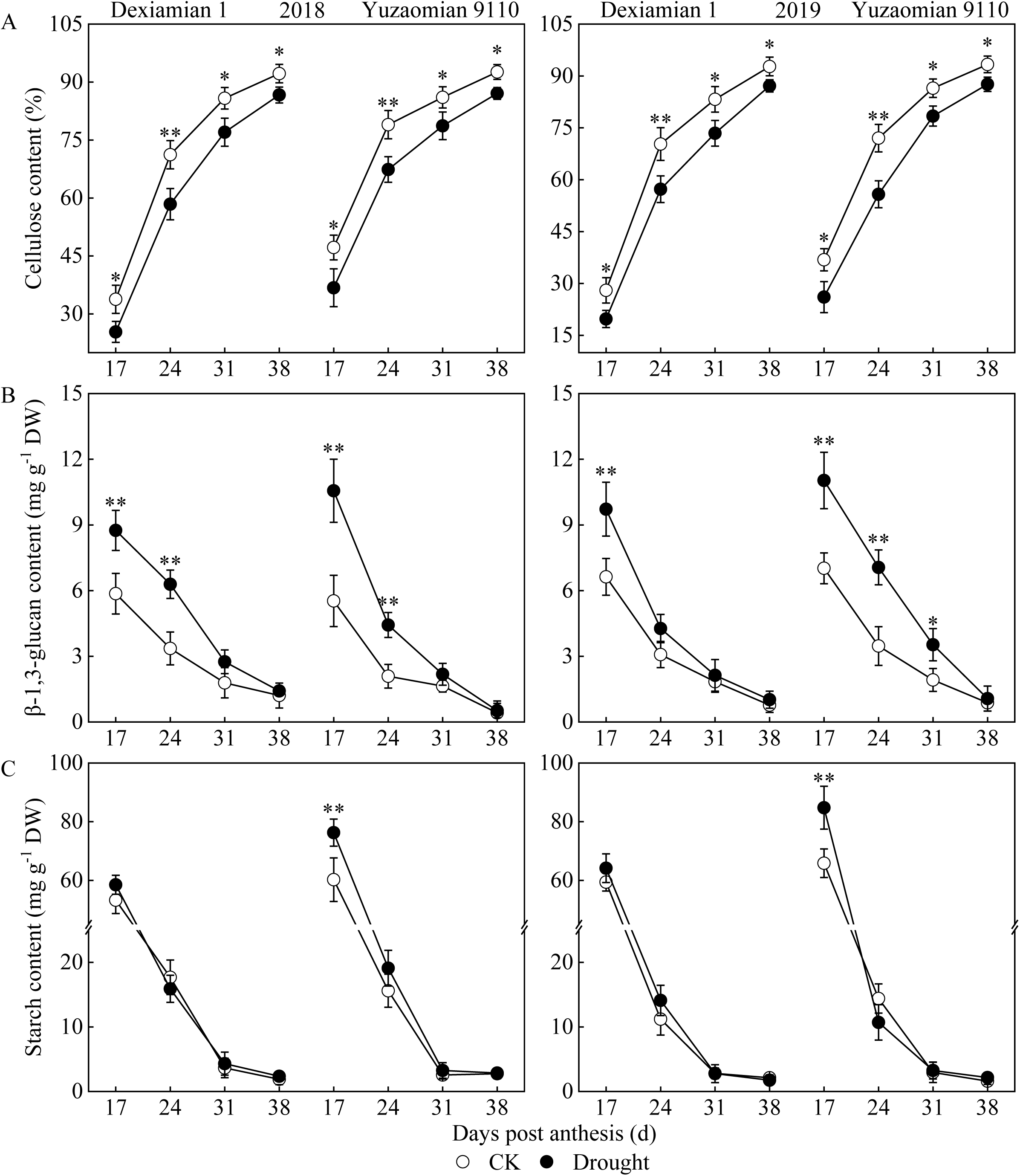
Effects of drought on the dynamic changes of cellulose content (A), β-1, 3-glucan content (B), and starch content (C) in cotton fiber in 2018 and 2019. Values are the means of three replicates ±SD. * and ** indicate statistically significant differences determined between different treatments at *P*=0.05 and *P*=0.01, respectively.

### Changes of glucose and fructose contents in cotton fiber

The contents of glucose and fructose in developing cotton fiber gradually decreased with DPA and reached extremely low levels at 38 DPA. Compared to CK, both glucose and fructose contents were significantly decreased during 17-24 DPA under drought (Fig. 6A, B). Taken together, drought induced huge physiological changes in the cotton fiber, especially at 17 DPA, which prompted us to further investigate the molecular responses.

**Fig. 6.**
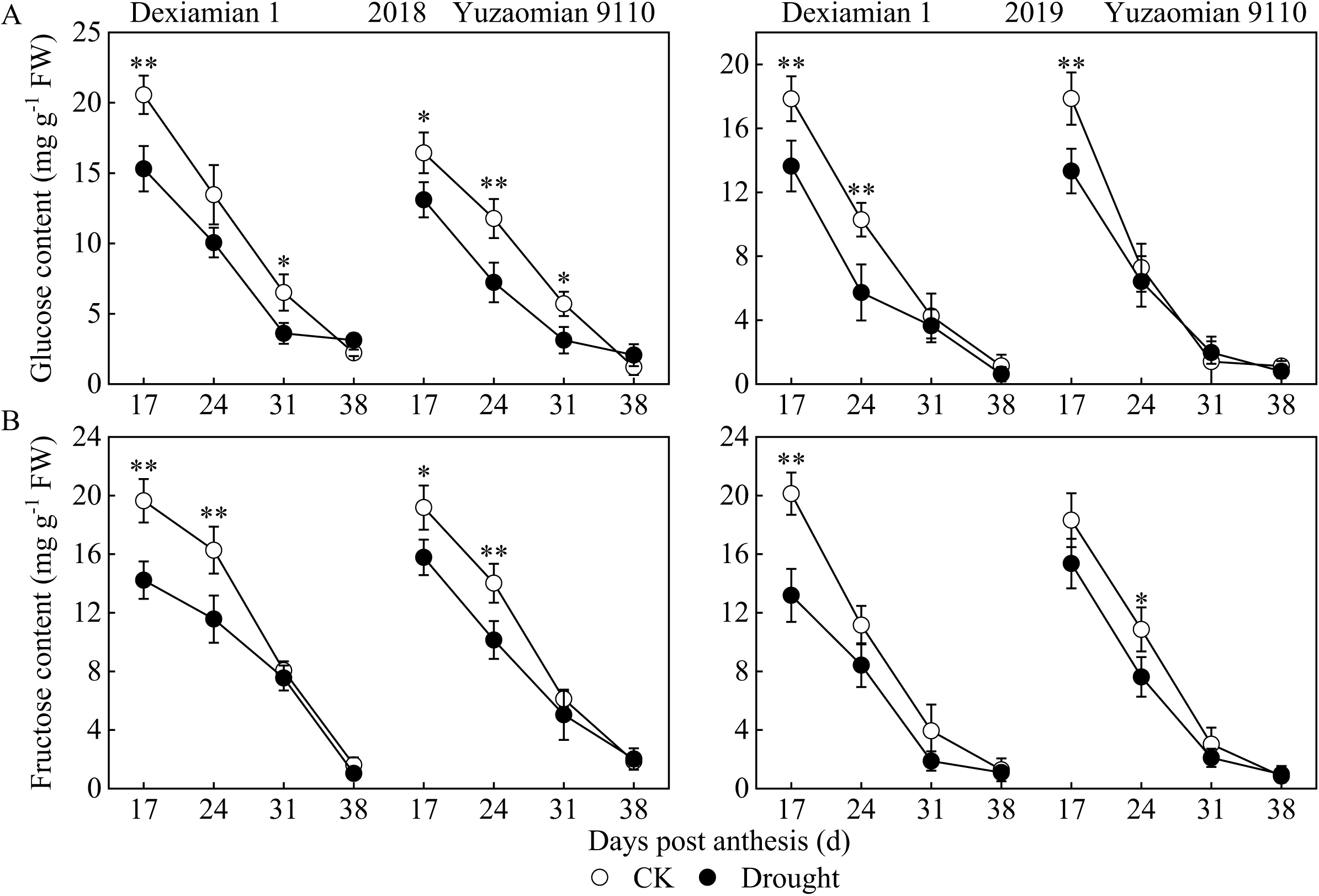
Effects of drought on the dynamic change of glucose content (A) and fructose content (B) in cotton fiber in 2018 and 2019. Values are the means of three replicates ±SD. * and ** indicate statistically significant differences determined between different treatments at *P*=0.05 and *P*=0.01, respectively.

### Differentially regulated gene networks in cotton fiber under drought

Cotton fibers at 17 DPA under CK and drought were sampled for transcriptional profile testing to identify drought-responsive genes that influenced secondary cell wall and fiber strength forming. The results revealed that differentially expressed genes (DEGs) induced by drought of 531 (274 up-regulated and 247 down-regulated) in Dexiamian 1 and 641 (458 up-regulated and 183 down-regulated) in Yuzaomian 9110 were detected respectively (Fig. 7A). Compared to CK, the majority of DEGs in cotton fiber at 17 DPA were up-regulated under drought. Moreover, 224 DEGs synchronously participated in the response to drought in two cultivars, accounting for 23.6% of total DEGs (Fig. 7B). However, most of them exhibited different changing modes in the two cultivars. For instance, genes involved in the oxidation-reduction process were up-regulated in Dexiamian 1 but down-regulated in Yuzaomian 9110, and the generation of ATP was suppressed in Dexiamian 1 but promoted in Yuzaomian 9110 (Fig. S3).

**Fig. 7.**
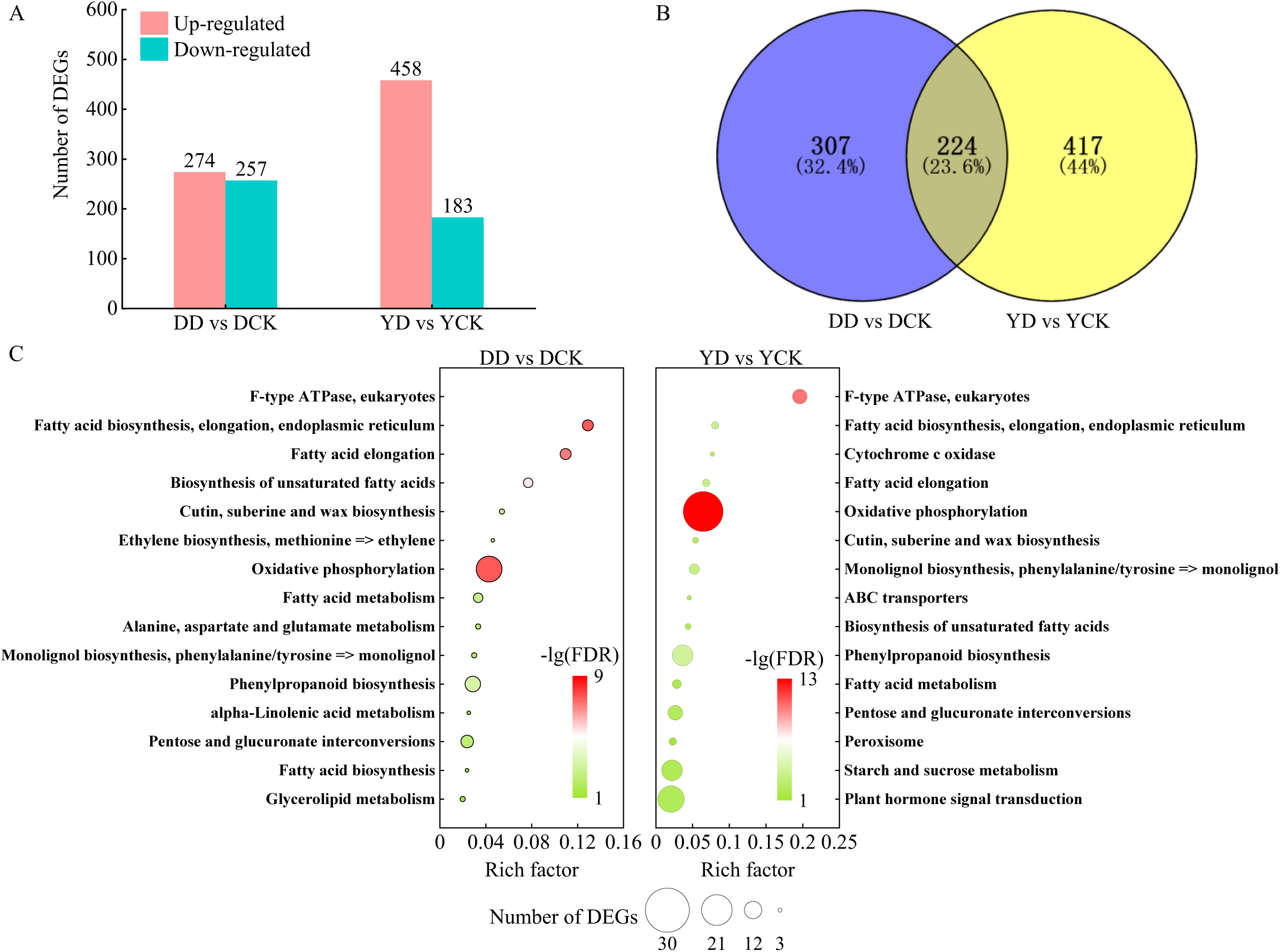
Number (A), venn diagram (B), and enriched KEGG pathways (C) of drought-induced differentially expressed genes (DEGs) in cotton fiber of Dexiamian 1 and Yuzaomian 9110. Fibers were sampled at 17 dasys post anthesis (DPA). DD and DCK respectively represent Dexiamian 1 under drought and control conditions, and YD and YCK respectively represent Yuzaomian 9110 under drought and control conditions. In Fig. 7C, rich factor refers to the ratio of the number of transcripts in the pathway entry in the differentially expressed transcript to the total number of transcripts in the transcript that are located in the pathway entry. The dot size shows the number of DEGs enriched in the pathway, and the dot color represents the -log10(FDR).

To further identify the drought-responsive metabolisms and pathways, the DEGs were annotated and classified by using both Gene Ontology (GO) analysis and Encyclopedia of Genes and Genomes (KEGG) analysis. The GO enrichment analysis results showed that the two cultivars shared many metabolisms in response to drought, like ‘response to oxidative stress’, ‘ATP metabolic process’, and ‘carbohydrate metabolic process’. Besides, ‘cellulose biosynthetic process’, ‘cellulular glucan metabolic process’, and ‘cell wall biogenesis’ were enriched in Dexiamian 1, and ‘cell wall modification’ and ‘cell wall organization’ were enriched in Yuzaomian 9110 (Fig. S4), which indicated the cell wall forming was strongly affected under drought. Regarding the KEGG analysis, the top 15 KEGG pathways were displayed in Fig. 7C. The analysis results revealed that pathways of ‘pentose and glucuronate interconversions’, ‘phenylpropanoid biosynthesis’, ‘fatty acid metabolism, and cutin’, ‘suberine and wax biosynthesis’ were significantly enriched in both cultivars under drought (Fig. 7C). The lignin metabolic pathway was assigned to ‘phenylpropanoid biosynthesis’ in both cultivars. Concerning the cellulose metabolic pathway, ‘starch and sucrose metabolism’ was enriched in Yuzaomian 9110 (Fig. 7C).

To further confirm the RNA-seq results, the expression levels of 15 randomly selected DEGs in cotton fiber at 17 DPA we determined with qRT-PCP for each cultivar. As shown in Fig. S5, the qRT-PCR results were well matched to the RNA-seq data with a high correlation coefficient (*R*=0.89), which validated the accuracy of the transcriptome.

### Expression patterns of genes related to sucrose metabolism and cell wall construction in cotton fiber under drought

During secondary wall thickening, cellulose and other cell wall components (β-1,3-glucan and lignin) are rapidly deposited in cotton fiber, and this requires an abundance of sucrose as substrates and energy supply. Thus the following analysis was focused on the DEGs related to sucrose metabolism and cell wall construction. As shown in Fig. 8, lignin synthesis, sucrose degrading, 1,3-glucan metabolism, glycolysis, and cellulose metabolism were notably regulated under drought.

**Fig. 8.**
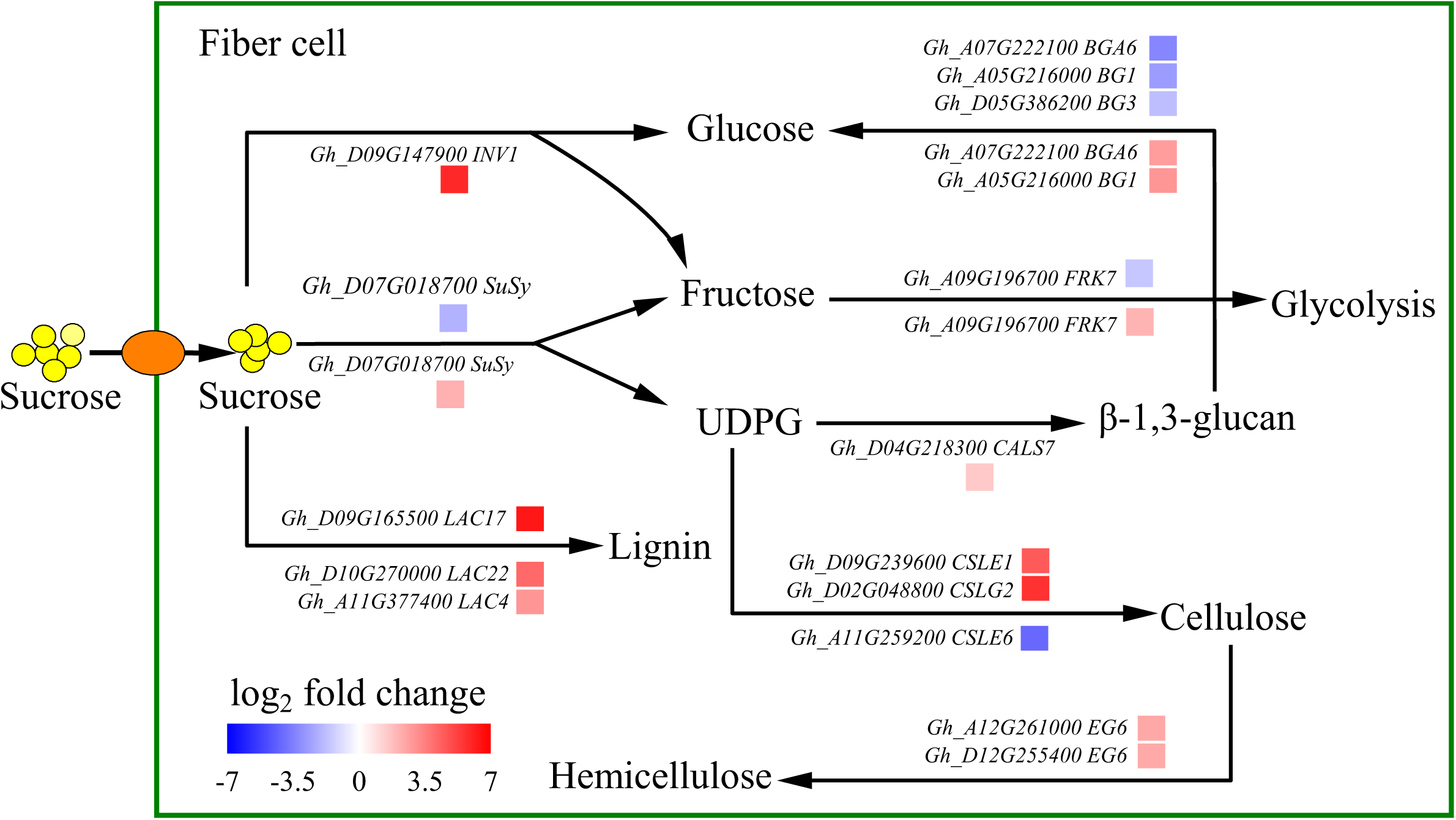
Drought-induced DEGs and pathways related to sucrose metabolism and cell wall construction in cotton fiber at 17 DPA. The box above and below the leader line indicate the varied genes in Dexiamian 1 and Yuzaomian 9110 under drought, respectively. The color scale represents the value of log2 fold change. UDPG, uridine-5’-diphosphoglucose.

In cotton fiber cells, the imported sucrose has to be converted into UDPG, glucose, and fructose by sucrose synthase (SuSy) and invertases (INV) before being directed into downstream metabolic processes like biosynthesis of β-1,3-glucan and cellulose, and glycolysis, etc. Under drought, the expression of *GhSuSy* was decreased in Dexiamian 1 while increased in Yuzaomian 9110, and *GhINV* was specifically up-regulated in Yuzaomian 9110 (Fig. 8, Fig. S6). Compared to CK, *GhFRK7* correlated with glycolysis exhibited increased expression in Dexiamian 1 but decreased expression in Yuzaomian 9110 under drought (Fig. 8, Fig. S6).

Consistently, the lignin biosynthesis pathway in cotton fiber was promoted in both cultivars under drought. Laccases (LACs) are key enzymes in lignin biosynthesis. Under drought, *GhLAC17* was up-regulated in Dexiamian 1, and *GhLAC4* and *GhLAC22* were up-regulated in Yuzaomian 9110 (Fig. 8, Fig. S6). Concerning β-1,3-glucan synthesis, *GhCALS7* that encodes a β-1,3-glucan synthesis-related protein was up-regulated in Yuzaomian 9110 under drought (Fig. 8, Fig. S6). In addition, β-1,3-glucanase (BG) genes associated with β-1,3-glucan degrading were down-regulated in Dexiamian 1 (*GhBGA6, GhBG1*, and *GhBG3*) and up-regulated in Yuzaomian 9110 (*GhBGA6* and *GhBG1*) under drought (Fig. 8, Fig. S6). Regarding the cellulose accumulation, two cellulose synthase-like protein genes (*GhCSLE1* and *GhCSLG2*) and two cellulose-degrading related genes encoding endoglucanase6 (EG6) were simultaneously up-regulated in Dexiamian 1 under drought. Whereas in Yuzaomian 9110, a cellulose synthase-like protein gene (*GhCLE6*) was down-regulated under drought (Fig. 8, Fig. S6).

## Discussion

### Drought decreased sucrose translocation from cottonseed coat to fiber

Drought is one of the most universal and destructive abiotic stress, which could do great harm to cotton production and fiber quality, especially fiber strength. Previous studies have reported that the occurrence of drought during the flowering and boll forming stage could lead to the biggest loss in fiber yield and quality (Snowden *et al*., 2014) including fiber strength (Wang *et al*., 2016; Gao *et al*., 2021). Consistent with the previous studies, the fiber strength was dramatically decreased under drought (Table. 1). Moreover, a significant difference in the interaction effect of water treatment (WT)×cultivar was observed in both 2018 and 2019 (Table. 1), since the drought-sensitive cultivar of Yuzaomian 9110 had a greater decreasing amplitude in fiber strength than the drought-tolerant cultivar of Dexiamian 1, which was also observed in a previous study (Basal *et al*., 2009), where STN-8A suffered a greater loss in fiber strength than Sahin-2000.

Cotton bolls are highly active sink organs during the flowering and boll forming stage, receiving photosynthates produced by source leaves (Pace *et al*., 1999). Previous studies found that although the photosynthetic rates of source leaves were decreased under drought (Pettigrew, 2004; Tsonev *et al*., 2011; Wang *et al*., 2016), the assimilate accumulation in source leaves was still promoted (Albacete *et al*., 2014). Lemoine *et al*. and Sevanto suggested that this was because the long-distance transport of photosynthates through the phloem was blocked (Lemoine *et al*., 2013; Sevanto, 2018). Hence, fewer photosynthates were transported to reproductive organs, resulting in the decreased boll weight (Wang *et al*., 2016). Within the boll, there are 26-30 cottonseeds and each cottonseed could be subdivided into three sink tissues: fiber, cottonseed coat, and embryo. In the present study, the final biomass and thickness of the cottonseed coat were little affected by drought, while the biomass of cotton fiber and embryo were significantly decreased (Table. S2, Fig. S2), indicating that lower boll weight due to drought reported by previous studies (Wang *et al*., 2016) was mainly reflected in the reduced biomass of fiber and embryo. The unaffected cottonseed coat might be an adaptive scheme to drought stress (Noodén *et al*., 1985). Moreover, results about the biomass distribution within cottonseed showed that the proportion of embryo biomass was little affected (except for Yuzaomian 9110 in 2019), and the proportion of cottonseed coat biomass was significantly increased, and the proportion of fiber biomass was dramatically decreased (Fig. 1A), indicating that the balance of biomass allocation within the boll was disturbed and the fiber was most adversely affected. Within the boll, the photosynthates are first delivered to cottonseed coat and then transport outwards to fiber and inwards to embryo (Ruan *et al*., 1997), so cottonseed coat as a nutrient transfer station plays a critical role in maintaining the balance of biomass allocation (Ruan and Furbank, 2003; Ruan, 2013). Compared with CK, the ^13^C distribution percentage (DP_13C_) was remarkably increased in cottonseed coat and was decreased in fiber under drought (Fig. 2), meaning that the carbon flow from cottonseed coat to fiber was impaired by drought, which explained why the proportion of cottonseed coat biomass was significantly increased, but the proportion of fiber biomass was dramatically decreased.

Sucrose is the main transport medium for carbon flow from cottonseed coat to fiber (Ruan *et al*., 1997), and the apoplasmic route for sucrose transport prevails at the later stage of fiber development when plasmodesmata are closed and sucrose transporter genes are highly expressed (Zhang *et al*., 2017). SUT (sucrose transporter) also called SUC (sucrose carrier), and SWEET (sugars will eventually be exported transporter) are two major identified sucrose transporter families in higher plants (Chen *et al*., 2015). Both proteins were readily detectable in the outer cottonseed (Ruan *et al*., 2000; Sun *et al*., 2019), and SUT proteins are responsible for the active uptake of sucrose from the apoplast against a concentration gradient (Zhou *et al*., 2007) while *GhSWEET10, GhSWEET12*, and *GhSWEET15* mediate sucrose efflux (Cox *et al*., 2017; Sun *et al*., 2019; Ding *et al*., 2021). In this study, drought notably enhanced the expressions of *GhSUT2A/D, GhSUT4A/D, GhSUT6A/D*, and *GhSUT8A/D* in the outer cottonseed coat at 17 DPA (Fig. 4), which facilitated the absorption of sucrose from the apoplast, resulting in higher sucrose concentration in cottonseed coat (Fig. 3A). A similar correlation between SUT gene expression and sucrose content in cottonseed coat was also reported in a previous study (Ding *et al*., 2019). However, under drought, the transcripts of *GhSWEET10* and *GhSWEET15* in the outer cottonseed coat were dramatically decreased at 17 DPA (Fig. 4), blocking sucrose outflow to fiber cells, which led to reduced sucrose content in the fiber (Fig. 3B). When the drought extended to 24 DPA, *GhSUT4A/D, GhSUT6A/D*, and *GhSUT8A/D* were significantly down-regulated, meaning that at this stage, the sucrose intake in seedcoat was suppressed. Similar results were obtained in developing soybean seeds as drought prolonged to the later developing stage (Zhao *et al*., 2020). However, compared to 17 DPA, more dramatic down-regulation of *GhSWEET10* and *GhSWEET15* at 24 DPA (Fig. 4) implied that the sucrose efflux from cottonseed coat to fiber was even more hampered leading to more sucrose remained in cottonseed coat, and this explained elevated sucrose level in cottonseed coat but decreased sucrose content in fiber (Fig. 3A, B). These results strongly indicated that under drought, the incoming sucrose in developing cottonseed was preferentially utilized by cottonseed coat to sustain its sink strength but not transferred to fiber, and this was probably a stress-adoptive strategy (Leisner *et al*., 2017).

### Drought reduced the flow of sucrose into cellulose synthesis within cotton fiber

Cellulose content in mature cotton fiber accounts for more than 90% (Haigler *et al*., 2012), and was reported to be positively correlated to fiber strength (Haigler *et al*., 2007). In cotton fiber, usually up to 80% of the incoming sucrose is partitioned to cellulose synthesis during the secondary wall synthesis stage (Haigler *et al*., 2001). Besides, sucrose is also the substrate for many other metabolisms in fiber, such as β-1,3-glucan (callose) synthesis, lignin synthesis, and starch synthesis, etc. and these metabolisms could affect cellulose synthesis and cell wall formation by competing for sucrose, especially under stresses (Velasco *et al*., 1994; Scheible and Pauly, 2004; Farrokhi *et al*., 2006; Bang *et al*., 2019). In the current study, transcriptome analysis by using fiber at 17 DPA showed that many drought-induced DEGs were closely correlated to lignin biosynthesis, accumulation of β-1,3-glucan and cellulose, and glycolysis (Fig. 8).

The sucrose degrading is essential for cotton fiber cell development since its cleavage products are the central molecules for cell wall construction, carbon partitioning, and energy generating (Haigler *et al*., 2001; Gou *et al*., 2007). Up to now, the known enzymatic paths of sucrose decomposition in plants are catalyzed by sucrose synthesis (SuSy) or invertases (INV) (Koch, 2004). SuSy reversibly converts sucrose into fructose and uridine-5’-diphosphoglucose (UDPG) (Braun *et al*., 2014), while INV irreversibly catalyzes sucrose into fructose and glucose (Roitsch and González, 2004). Under drought, the expression of *GhSuSy* was up-regulated in Yuzaomian 9110 and down-regulated in Dexiamian 1 (Fig. 8, Fig. S6), meaning that the production of UDPG was promoted in Yuzaomian 9110 and restricted in Dexiamian 1. In addition, up-regulating SuSy gene contributed to increased starch level (Muñoz *et al*., 2005), and this explained the elevated starch content in Yuzaomian 9110 under drought (Fig. 5C). The transcript level of *GhINV1* under drought was specifically enhanced in Yuzaomian 9110 (Fig. 8, Fig. S6) implying promoted production of hexose (glucose and fructose), which was probably intended to increase carbon availability or to strengthen osmotic adjustment to improve drought resistance(Kim *et al*., 2000; Yang *et al*., 2004).

In Yuzaomian 9110, the transcript of *GhFRK7* encoding a fructokinase that mediates fructose phosphorylation was significantly increased under drought (Fig. 8, Fig. S6), which prevented fructose accumulation (Fig. 6). In addition, fructokinase is a core kinase in glycolysis pathway (Plaxton, 1996). In *Hevea brasiliensis*, increased transcript abundance of *HbFRK2* gene strengthened glycolysis and energy generating (Fang *et al*., 2021). Similarly, *GhFRK7* and *GhATPA* showed higher expression levels in Yuzaomian 9110 under drought (Fig. 8, Fig. S3, Fig. S6), indicating promoted glycolysis and ATP generating, which probably contributed to meeting with higher energy demand in maintaining homeostasis under drought (Velasco *et al*., 1994). And as a result, more sucrose being directed into glycolysis and energy producing, the sucrose available to cell wall synthesis was decreased under drought. Similarly, reduced kernel weight in wheat was observed with up-regulation of glycolysis enzymes under heat stress (Rollins *et al*., 2013).

UDPG, the decomposition product of sucrose by SuSy, is the precursor to the synthesis of β-1, 3-glucan and cellulose, which is competed by these two metabolisms (Amor *et al*., 1995). β-1,3-glucan commonly regarded as callose in higher plants was synthesized by callose synthases (CALS) in a small amount during the early stage of fiber thickening development and was decomposed by β-1,3-glucanases (BG) at the later stage (Maltby *et al*., 1979; Tokumoto *et al*., 2002). For Yuzaomian 9110 under drought, the expression of *GhCALS7* was increased (Fig. 8, Fig. S6), and this indicated more UDPG was directed into β-1,3-glucan synthesis. Whereas in Dexiamian 1, no β-1,3-glucan synthesis-related gene was induced under drought, however, the β-1,3-glucan degrading process was significantly repressed by down-regulating *GhBGA6, GhBG1*, and *GHBG3* (Fig. 8, Fig. S6), facilitating the β-1,3-glucan deposition. Therefore, increased β-1,3-glucan content was observed in both cultivars under drought (Fig. 5B), which was consistent with the previous study (Gao *et al*., 2020).

Lignin is complex phenylpropanoid polymers (Chen *et al*., 2012). Although the lignin exists in cotton fiber with a small amount, it has significant functions in secondary wall synthesis and fiber quality formation (Gao *et al*., 2019). And its biosynthesis is inseparable from the carbon skeleton provided by sucrose (Amthor, 2003). Hence, the deposition of lignin and cellulose could be adjusted in a compensatory way. For example, Hu *et al*. reported the lignin content was reduced by up to 45% by suppressing a lignin biosynthesis related gene, however, a 15% increase in cellulose compensated for this reduction (Hu *et al*., 1999). Emerging studies showed that drought could promote lignin synthesis to enhance drought resistance in rice (*Oryza sativa*), oriental melon (*Cucumis melo*), and grapevine (*Vitis vinifera*) (Bang *et al*., 2019; Liu *et al*., 2020; Tu *et al*., 2020). Without exception, our data showed that ‘phenylpropanoid biosynthesis’ that is closely associated with lignin synthesis was significantly regulated in developing cotton fiber under drought (Fig. 7C). Laccases (LACs) are crucial for lignin biosynthesis, and the higher enzymatic activity of laccases corresponded to higher lignin content in cotton fiber (Balasubramanian *et al*., 2016). Under drought, the transcripts of *GhLAC4, GhLAC17*, and *GhLAC22* in cotton fiber were significantly increased (Fig. 8, Fig. S6), suggesting that lignin synthesis process would be promoted, which might reduce sucrose partitioned to cellulose synthesis to some extent. Hence, decreased cellulose content was observed under drought (Fig. 5A).

From the foregoing, drought increased sucrose flow to starch, glycolysis, β-1,3-glucan, and lignin, and this meant reduced sucrose flux to cellulose synthesis in the case of limited sucrose. Moreover, a cellulose synthase-like protein gene (*GhCSLE*6) was down-regulated in Yuzaomian 9110 under drought (Fig. 8, Fig. S6), directly limiting the cellulose synthesis. Whereas in Dexiamina 1, two cellulose synthase-like protein genes (*GhCSLG2* and *GhCSLE1*) displayed relatively higher expression levels under drought (Fig. 8, Fig. S6), which seemed to imply promoted cellulose synthesis. Indeed, expressions of genes (*GhEG6*) encoding endoglucanase that converts the cellulose into hemicellulose (Moneo-Sánchez *et al*., 2020) were significantly increased in Dexiamian 1 under drought (Fig. 8, Fig. S6), which could restrict cellulose accumulation since cellulose deposit is controlled by both cellulose synthesis and degradation processes (Gou *et al*., 2007). Therefore, both cultivars exhibited reduced cellulose content under drought (Fig. 5A).

## Conclusion

Drought altered the biomass distribution modes within the cottonseed: the proportion of biomass partitioned to cottonseed coat was significantly increased, while the proportion distributed to fiber was dramatically decreased. And this was largely due to blocked photosynthates (sucrose) translocation from the cottonseed coat to the fiber under drought. Further, the down-regulation of *GhSWEET10* and *GhSWEET15* in the outer cottonseed coat was responsible for the restricted sucrose transfer under drought. In cotton fiber, lignin synthesis and β-1,3-glucan deposition were promoted under drought, leading to decreased sucrose flux to cellulose synthesis. Additionally, starch synthesis and glycolysis were exclusively enhanced in Yuzaomian 9110 under drought, which further reduced sucrose flow into cellulose. Under drought, the cellulose synthesis was stunted due to down-regulation of cellulose synthase-like protein gene (*GhCSLE*6) in Yuzaomian 1, while the cellulose deposition was impaired due to promoted cellulose degrading process in Dexiamian 1. In brief, blocked sucrose translocation from cottonseed coat to fiber and reduced sucrose partitioned to cellulose resulted in reduced fiber cellulose content and inferior fiber strength under drought. Overall, this study revealed novel insights into understanding how drought affected cotton fiber strength and provided new ideas on cotton breeding for resistance against drought.

## Abbreviations

DEGs: differentially expressed genes
DPA: days post anthesis
INV: invertase
SuSy: sucrose synthetase
SUT: sucrose transport
SWEET: sugars will eventually be exported transporter
UDPG: uridine-5’-diphosphoglucose

## Supplementary data

Table S1. Primers used for qRT-PCR in this study.

Table S2. Effects of drought on the biomass of embryo, cottonseed coat, and fiber per seed at maturity.

Fig. S1. Effects of drought on the accumulation of cotton fiber biomass per seed.

Fig. S2. Effects of drought on the dynamic changes of thickness of cottonseed coat, inner cottonseed coat, and outer cottonseed coat.

Fig. S3. Comparison of drought-induced differentially expressed genes (DEGs) involved in oxidation-reduction process and ATP metabolic process in cotton fiber at 17 DPA of Dexiamian 1 and Yuzaomian 9110.

Fig. S4. Enriched GO terms of drought-induced DEGs in cotton fiber at 17 DPA of Dexiamian 1 and Yuzaomian 9110.

Fig. S5. Comparison of the relative expression values of randomly selected drought-induced DEGs in cotton fiber at 17 DPA determined by RNA-seq and qRT-PCR.

Fig. S6. The relative expression levels of selected drought-responsive genes related to sucrose metabolism and cell wall construction by qRT-PCR in Dexiamian 1 and Yuzaomian 9110

## Acknowledgment

This work was supported by the National Natural Science Foundation of China (31630051, 31901463), China Agriculture Research System of MOF and MARA (CARS-15-14), Natural Science Foundation of Jiangsu Province (BK20190524), the China Postdoctoral Science Foundation (2020M681633), Jiangsu Collaborative Innovation Center for Modern Crop Production (JCIC-MCP) and High-level Talent Introduction Program of Nanjing Agricultural University.

## Author contribution

Zhiguo Zhou, Wei Hu, Youhua Wang, Yali Meng, Binglin Chen, Wenqing Zhao, and Shanshan Wang conceived the project and designed the experiments. Honghai Zhu, Yuxia Li, Jie Zou, and Jiaqi He performed the experiments, and the data were analyzed by Honghai Zhu, Yuxia Li, and Jie Zou. The manuscript was written by Honghai Zhu and Wei Hu.

## Notes

### Competing Interest Statement

The authors have declared no competing interest.

## References

Abdelraheem A, Esmaeili N, O’Connell M, Zhang JF. 2019. Progress and perspective on drought and salt stress tolerance in cotton. Industrial Crops and Products 130, 118–129.

Albacete AA, Martínez-Andújar C, Pérez-Alfocea F. 2014. Hormonal and metabolic regulation of source– sink relations under salinity and drought: from plant survival to crop yield stability. Biotechnology Advances 32, 12–30.

Amor Y, Haigler CH, Johnson S, Wainscott M, Delmer DP. 1995. A membrane-associated form of sucrose synthase and its potential role in synthesis of cellulose and callose in plants. Proceedings of the National Academy of Sciences of the United States of America 92, 9353–9357.

Amthor JS. 2003. Efficiency of lignin biosynthesis: a quantitative analysis. Annals of Botany 91, 673–695.

Ayre BG. 2011. Membrane-transport systems for sucrose in relation to whole-plant carbon partitioning. Molecular Plant 4, 377–394.

Baker RF, Leach KA, Braun DM. 2012. SWEET as sugar: new sucrose effluxers in plants. Molecular Plant 5, 766–768.

Balasubramanian VK, Rai KM, Thu SW, Hii MM, Mendu V. 2016. Genome-wide identification of multifunctional laccase gene family in cotton (Gossypium spp.); expression and biochemical analysis during fiber development. Scientific Reports 6, 34309.

Bang SW, Lee DK, Jung H, Chung PJ, Kim YS, Choi YD, Suh JW, Kim JK. 2019. Overexpression of OsTF1L, a rice HD-Zip transcription factor, promotes lignin biosynthesis and stomatal closure that improves drought tolerance. Plant Biotechnology Journal 17, 118–131.

Basal H, Dagdelen N, Unay A, Yilmaz E. 2009. Effects of deficit drip irrigation ratios on cotton (Gossypium hirsutum L.) yield and fibre quality. Journal of Agronomy and Crop Science 195, 19–29.

Braun DM, Wang L, Ruan YL. 2014. Understanding and manipulating sucrose phloem loading, unloading, metabolism, and signalling to enhance crop yield and food security. Journal of Experimental Botany 65, 1713–1735.

Carpita NC, Delmer DP. 1981. Concentration and metabolic turnover of UDP-glucose in developing cotton fibers. Journal of Biological Chemistry 256, 308–315.

Chen F, Tobimatsu Y, Havkin-Frenkel D, Dixon RA, Ralph J. 2012. A polymer of caffeyl alcohol in plant seeds. Proceedings of the National Academy of Sciences of the United States of America 109, 1772–1777.

Chen LQ, Cheung LS, Feng L, Tanner W, Frommer WB. 2015. Transport of sugars. Annual Review of Biochemistry 84, 865–894.

Coleman HD, Yan J, Mansfield SD. 2009. Sucrose synthase affects carbon partitioning to increase cellulose production and altered cell wall ultrastructure. Proceedings of the National Academy of Sciences of the United States of America 106, 13118–13123.

Cox KL, Meng FH, Wilkins KE, Li FJ, Wang P, Booher NJ, Carpenter SCD, Chen LQ, Zheng H, Gao XQ, et al. 2017. TAL effector driven induction of a SWEET gene confers susceptibility to bacterial blight of cotton. Nature Communications 8, 15588.

Dağdelen N, Başal H, Yılmaz E, Gürbüz T, Akçay S. 2009. Different drip irrigation regimes affect cotton yield, water use efficiency and fiber quality in western Turkey. Agricultural Water Management 96, 111–120.

Dai AG. 2011. Drought under global warming: a review. Wiley Interdisciplinary Reviews: Climate Change 2, 45–65.

De Schepper V, De Swaef T, Bauweraerts I, Steppe K. 2013. Phloem transport: a review of mechanisms and controls. Journal of Experimental Botany 64, 4839–4850.

Ding XY, Li XB, Wang L, Zeng JY, Huang L, Xiong L, Song SQ, Zhao J, Hou L, Wang FL, et al. 2021. Sucrose-enhanced reactive oxygen species generation promotes cotton fiber initiation and secondary cell wall deposition. Plant Biotechnology Journal 6, 1092–1094.

Ding XY, Zeng JY, Huang L, Li XB, Song SQ, Pei Y. 2019. Senescence-induced expression of ZmSUT1 in cotton delays leaf senescence while the seed coat-specific expression increases yield. Plant Cell Reports 38, 991–1000.

Fang PC, Long XY, Fang YJ, Chen H, Yu M. 2021. A predominant isoform of fructokinase, HbFRK2, is involved in Hevea brasiliensis (para rubber tree) latex yield and regeneration. Plant Physiology and Biochemistry 162, 211–220.

Farrokhi N, Burton RA, Brownfield L, Hrmova M, Wilson SM, Bacic A, Fincher GB. 2006. Plant cell wall biosynthesis: genetic, biochemical and functional genomics approaches to the identification of key genes. Plant Biotechnology Journal 4, 145–167.

Gao M, Snider JL, Bai H, Hu W, Wang R, Meng YL, Wang YH, Chen BL, Zhou ZG. 2020. Drought effects on cotton (Gossypium hirsutum L.) fibre quality and fibre sucrose metabolism during the flowering and boll-formation period. Journal of Agronomy and Crop Science 206, 309–321.

Gao M, Xu BJ, Wang YH, Zhou ZG, Hu W. 2021. Quantifying individual and interactive effects of elevated temperature and drought stress on cotton yield and fibre quality. Journal of Agronomy and Crop Science 207, 422–436.

Gao ZY, Sun WJ, Wang J, Zhao CY, Zuo KJ. 2019. GhbHLH18 negatively regulates fiber strength and length by enhancing lignin biosynthesis in cotton fibers. Plant Science 286, 7–16.

Gou JY, Wang LJ, Chen SP, Hu WL, Chen XY. 2007. Gene expression and metabolite profiles of cotton fiber during cell elongation and secondary cell wall synthesis. Cell Research 17, 422–434.

Haigler CH, Lissete B, Stiff MR, Tuttle JR. 2012. Cotton fiber: a powerful single-cell model for cell wall and cellulose research. Frontiers in Plant Science 3, 104.

Haigler CH, Singh B, Zhang DS, Hwang SJ, Wu CF, Cai WX, Hozain M, Kang W, Kiedaisch B, Strauss RE. 2007. Transgenic cotton over-producing spinach sucrose phosphate synthase showed enhanced leaf sucrose synthesis and improved fiber quality under controlled environmental conditions. Plant Molecular Biology 63, 815–832.

Haigler CH, Ivanova-Datcheva M, Hogan PS, Salnikov VV, Hwang S, Martin K, Delmer DP. 2001. Carbon partitioning to cellulose synthesis. Plant Molecular Biology 47, 29–51.

Harrison CJ, Hedley CL, Wang TL. 1998. Evidence that the rug3 locus of pea (Pisum sativum L.) encodes plastidial phosphoglucomutase confirms that the imported substrate for starch synthesis in pea amyloplasts is glucose-6-phosphate. The Plant Journal 13, 753–762.

Hendrix DL. 1990. Carbohydrates and carbohydrate enzymes in developing cotton ovules. Physiologia Plantarum 78, 85–92.

Hu W, Snider JL, Wang HM, Zhou ZG, Chastain DR, Whitaker J, Perry CD, Bourland FM. 2018. Water-induced variation in yield and quality can be explained by altered yield component contributions in field-grown cotton. Field Crops Research 224, 139–147.

Hu W, Cao YT, Loka DA, Harris-Shultz KR, Reiter RJ, Ali S, Liu Y, Zhou ZG. 2020. Exogenous melatonin improves cotton (Gossypium hirsutum L.) pollen fertility under drought by regulating carbohydrate metabolism in male tissues. Plant physiology and biochemistry 151, 579–588.

Hu W, Loka DA, Fitzsimons TR, Zhou ZG, Oosterhuis DM. 2018. Potassium deficiency limits reproductive success by altering carbohydrate and protein balances in cotton (Gossypium hirsutum L.). Environmental and Experimental Botany 145, 87–94.

Hu WJ, Harding SA, Lung J, Popko JL, Ralph J, Stokke DD, Tsai CJ, Chiang VL. 1999. Repression of lignin biosynthesis promotes cellulose accumulation and growth in transgenic trees. Nature Biotechnology 17, 808–812.

Huang G, Huang JQ, Chen XY, Zhu YX. 2021. Recent advances and future perspectives in cotton research. Annual Review of Plant Biology 72, 437–462.

Ibrahim W, Zhu YM, Chen Y, Qiu CW, Zhu SJ, Wang FB. 2019. Genotypic differences in leaf secondary metabolism, plant hormones and yield under alone and combined stress of drought and salinity in cotton genotypes. Physiologia Plantarum 165, 343–355.

Kanehisa M, Araki M, Goto S, Hattori M, Hirakawa M, Itoh M, Katayama T, Kawashima S, Okuda S, Tokimatsu T, et al. 2008. KEGG for linking genomes to life and the environment. Nucleic Acids Research 36, D480–D484.

Kim JY, Mahé A, Brangeon J, Prioul JL. 2000. A maize vacuolar invertase, IVR2, is induced by water stress. Organ/tissue specificity and diurnal modulation of expression. Plant physiology 124, 71–84.

Koch K. 2004. Sucrose metabolism: regulatory mechanisms and pivotal roles in sugar sensing and plant development. Current Opinion in Plant Biology 7, 235–246.

Köhle H, Jeblick W, Poten F, Blaschek W, Kauss H. 1985. Chitosan-elicited callose synthesis in soybean cells as a ca-dependent process. Plant Physiology 77, 544–551.

Kühn C, Grof CP. 2010. Sucrose transporters of higher plants. Current Opinion in Plant Biology 13, 288–298.

Leisner CP, Yendrek CR, Ainsworth EA. 2017. Physiological and transcriptomic responses in the seed coat of field-grown soybean (Glycine max L. Merr.) to abiotic stress. BMC Plant Biology 17, 242.

Li W, Sun K, Ren ZY, Song CX, Pei XY, Liu YA, Wang ZY, He KL, Zhang F, Zhou XJ, et al. 2018. Molecular evolution and stress and phytohormone responsiveness of SUT genes in Gossypium hirsutum. Frontiers in Genetics 9, 494.

Liu W, Jiang Y, Wang CH, Zhao LL, Jin YZ, Xing QJ, Li M, Lv TH, Qi HY. 2020. Lignin synthesized by CmCAD2 and CmCAD3 in oriental melon (Cucumis melo L.) seedlings contributes to drought tolerance. Plant Molecular Biology 103, 689–704.

Maltby D, Carpita NC, Montezinos D, Kulow C, Delmer DP. 1979. β-1,3-glucan in developing cotton fibers. Plant Physiology 63, 42–44.

Moneo-Sánchez M, Vaquero-Rodríguez A, Hernández-Nistal J, Albornos L, Knox P, Dopico B, Labrador E, Martín I. 2020. Pectic galactan affects cell wall architecture during secondary cell wall deposition. Planta 251, 100.

Muñoz FJ, Baroja-Fernández E, Morán-Zorzano MT, Viale AM, Etxeberria E, Alonso-Casajús N, Pozueta-Romero J. 2005. Sucrose synthase controls both intracellular ADP glucose levels and transitory starch biosynthesis in source leaves. Plant Cell Physiology 46, 1366–1376.

Noodén LD, Blakley KA, Grzybowski JM. 1985. Control of seed coat thickness and permeability in soybean: a possible adaptation to stress. Plant Physiology 79, 543–545.

Pace PF, Cralle HT, Cothren JT, Senseman SA. 1999. Photosynthate and dry matter partitioning in short-and long-season cotton cultivars. Crop Science 39, 1065–1069.

Peng L, Hocart CH, Redmond JW, Williamson RE. 2000. Fractionation of carbohydrates in Arabidopsis root cell walls shows that three radial swelling loci are specifically involved in cellulose production. Planta 211, 406–414.

Peng Q, Cai YM, Lai EH, Nakamura M, Liao L, Zheng BB, Ogutu C, Cherono S, Han YP. 2020. The sucrose transporter MdSUT4.1 participates in the regulation of fruit sugar accumulation in apple. BMC Plant Biology 20, 191.

Pettigrew WT. 2004. Physiological consequences of moisture deficit stress in cotton. Crop Science 44, 1265–1272.

Plaxton WC. 1996. The organization and regulation of plant glycolysis. Annual Review of Plant Physiology and Plant Molecular Biology 47, 185–214.

Pugh DA, Offler CE, Talbot MJ, Ruan YL. 2010. Evidence for the role of transfer cells in the evolutionary increase in seed and fiber biomass yield in cotton. Molecular Plant 3, 1075–1086.

Read SM, Bacic T. 2002. Plant biology. Prime time for cellulose. Science 295, 59–60.

Roitsch T, González M. 2004. Function and regulation of plant invertases: sweet sensations. Trends in Plant Science 9, 606–613.

Rollins JA, Habte E, Templer S, Colby T, Schmidt J, Korff M. 2013. Leaf proteome alterations in the context of physiological and morphological responses to drought and heat stress in barley (Hordeum vulgare L.). Journal of Experimental Botany 64, 3201–3212.

Ruan YL. 2013. Boosting seed development as a new strategy to increase cotton fiber yield and quality. Journal of Integrative Plant Biology 55, 572–575.

Ruan YL, Chourey PS, Delmer DP, Perezgrau L. 1997. The differential expression of sucrose synthase in relation to diverse patterns of carbon partitioning in developing cotton seed. Plant Physiology 115, 375–385.

Ruan YL, Furbank RT. 2003. Suppression of sucrose synthase gene expression represses cotton fiber cell initiation, elongation, and seed development. The Plant Cell 15, 952–964.

Ruan YL, Chourey PS. 1998. A fiberless seed mutation in cotton is associated with lack of fiber cell initiation in ovule epidermis and alterations in sucrose synthase expression and carbon partitioning in developing seeds. Plant Physiology 118, 399–406.

Ruan YL, Llewellyn DJ, Furbank RT. 2000. Pathway and control of sucrose import into initiating cotton fibre cells. Functional Plant Biology 27, 795–800.

Ruehr NK, Offermann CA, Gessler A, Winkler JB, Ferrio JP, Buchmann N, Barnard RL. 2009. Drought effects on allocation of recent carbon: from beech leaves to soil CO2 efflux. New Phytologist 184, 950–961.

Scheible WR, Pauly M. 2004. Glycosyltransferases and cell wall biosynthesis: novel players and insights. Current Opinion in Plant Biology 7, 285–295.

Snowden MC, Ritchie GL, Simao FR, Bordovsky JP. 2014. Timing of episodic drought can be critical in cotton. Agronomy Journal 106, 452–458.

Sun WJ, Gao ZY, Wang J, Huang YQ, Chen Y, Li JF, Lv ML, Wang J, Luo M, Zuo KJ. 2019. Cotton fiber elongation requires the transcription factor GhMYB212 to regulate sucrose transportation into expanding fibers. New Phytologist 222, 864–881.

Tokumoto H, Wakabayashi K, Kamisaka S, Hoson T. 2002. Changes in the sugar composition and molecular mass distribution of matrix polysaccharides during cotton fiber development. Plant Cell Physiology 43, 411–418.

Tsonev T, Velikova V, Yildiz-Aktas L, Gürel A, Edreva A. 2011. Effect of water deficit and potassium fertilization on photosynthetic activity in cotton plants. Plant Biosystems 145, 841–847.

Tu MX, Wang XH, Yin WC, Wang Y, Li YJ, Zhang GF, Li Z, Song JY, Wang XP. 2020. Grapevine VlbZIP30 improves drought resistance by directly activating VvNAC17 and promoting lignin biosynthesis through the regulation of three peroxidase genes. Horticulture Research 7, 150.

Updegraff DM. 1969. Semimicro determination of cellulose inbiological materials. Analytical Biochemistry 32, 420–424.

Velasco R, Salamini F, Bartels D. 1994. Dehydration and ABA increase mRNA levels and enzyme activity of cytosolic GAPDH in the resurrection plant Craterostigma plantagineum. Plant Molecular Biology 26, 541–546.

Vera AR, Pongor LS, Mariño-Ramírez L, Landsman D. 2019. TPMCalculator: one-step software to quantify mRNA abundance of genomic features. Bioinformatics 35, 1960–1962.

Viglioni P, Verhalen LM, Kirkham MB, Mcnew RW. 1998. Screening cotton genotypes for seedling drought tolerance. Genetics & Molecular Biology 21, 545–549.

Wang R, Ji S, Zhang P, Meng YL, Wang YH, Chen BL, Zhou ZG. 2016. Drought effects on cotton yield and fiber quality on different fruiting branches. Crop Science 56, 1265–1276.

Wang R, Gao M, Ji S, Wang SS, Meng YL, Zhou ZG. 2016. Carbon allocation, osmotic adjustment, antioxidant capacity and growth in cotton under long-term soil drought during flowering and boll-forming period. Plant Physiology and Biochemistry 107, 137–146.

Yang J, Zhang J, Wang Z, Xu G, Zhu Q. 2004. Activities of key enzymes in sucrose-to-starch conversion in wheat grains subjected to water deficit during grain filling. Plant Physiology 135, 1621–1629.

Young MD, Wakefield MJ, Smyth GK, Oshlack A. 2010. Gene ontology analysis for RNA-seq: accounting for selection bias. Genome Biology 11, R14.

Zhang CK, Turgeon R. 2018. Mechanisms of phloem loading. Current Opinion in Plant Biology 43, 71–75.

Zhang ZY, Ruan YL, Zhou N, Wang F, Guan XY, Fang L, Shang XG, Guo WZ, Zhu SJ, Zhang TZ. 2017. Suppressing a putative sterol carrier gene reduces plasmodesmal permeability and activates sucrose transporter genes during cotton fiber elongation. The Plant Cell 29, 2027–2046.

Zhao Q, Chen L, Yao X, Zhang H, Wu J, Xie F. 2020. Effect of drought stress during soybean R2-R6 growth stages on sucrose metabolism in leaf and seed. International Journal of Molecular Sciences 21, 618.

Zhou YC, Qu HX, Dibley KE, Offler CE, Patrick JW. 2007. A suite of sucrose transporters expressed in coats of developing legume seeds includes novel pH-independent facilitators. Plant Journal 49, 750–764.

Zou J, Hu W, Li YX, He JQ, Zhu HH, Zhou ZG. 2020. Screening of drought resistance indices and evaluation of drought resistance in cotton (Gossypium hirsutum L.). Journal of Integrative Agriculture 19, 495–508.

